# Resolving the fate of trace metals during microbial remineralization of phytoplankton biomass in precursor banded iron formation sediments

**DOI:** 10.1101/2022.06.14.496128

**Authors:** Kathryn Rico, Manuel Schad, Aude Picard, Andreas Kappler, Kurt Konhauser, Nagissa Mahmoudi

## Abstract

Banded Iron Formations (BIFs) have long been considered a sedimentary record of seawater trace metal composition during the Precambrian. However, recent work has suggested that the trace metal composition of BIFs was derived from phytoplankton biomass, not seawater. In this model, phytoplankton biomass settles from the photic zone to the seafloor sediments, where it is then oxidized by heterotrophic microbes, such as dissimilatory Fe(III) reducing (DIR) bacteria, for energy generation. Remineralization of this biomass released the trace metals associated with organic molecules from phytoplankton (*i*.*e*., in metalloproteins), allowing these metals to be captured by Fe (oxyhydr)oxides and preserved in BIFs. While there is compelling evidence that the phytoplankton biomass served as a trace metal shuttle to precursor BIF sediments, it is unclear whether the degradation of biomass by DIR bacteria would liberate the biogenic trace metals as the model proposes. This work tests this hypothesis by using anoxic incubations of a model DIR bacterium (*Shewanella oneidensis* MR-1) with phytoplankton biomass as energy and carbon sources and ferrihydrite, a poorly crystalline Fe(III) oxyhydroxide (Fe(OH)_3_), as electron acceptor. Our results show that while *S. oneidensis* MR-1 can consume some of the carbon substrates found in phytoplankton biomass, there is no evidence that *S. oneidensis* MR-1 degraded metalloproteins which would have liberated trace metals. In the context of the Precambrian, these data imply that other heterotrophic bacteria, such as fermenters, may have had a larger role in the liberation of trace metals from dead biomass during early BIF development.

**Highlights:**

1. Phytoplankton are the proposed source of trace metals to banded iron formations
2. Iron reducers are hypothesized to release metals from phytoplankton biomass
3. Experiments show that iron reducers do not liberate metals when degrading biomass
4. Other microbial heterotrophs must have liberated the biogenic trace metals

## 1. Introduction

All life requires bioessential trace metals that act as structural components and reactive centers in metalloproteins (Jelen et al., 2016). Trends in trace metal concentrations in the sedimentary record through time reflect not only their availability in the oceans, but also their potential influence on the emergence or expansion of certain metabolisms (Robbins et al., 2016, and references therein). Over the course of Earth’s history, shifts in global redox state have greatly influenced the availability of trace metals in the ocean (*e*.*g*., Swanner et al., 2014). This is particularly true during the Proterozoic (2.5–0.5 Ga), where Earth’s atmosphere and oceans gradually oxygenated, microbial life diversified, and eukaryotic life evolved (Lyons et al., 2014).

Our understanding of trace metal contents in ancient oceans comes from various sedimentary deposits such as banded iron formations (BIFs), black shales, and authigenic pyrites (Robbins et al., 2016). Of these, BIFs are unique in that they precipitated directly out of seawater throughout the Precambrian, making them particularly useful archives to track secular changes in trace metal composition, and ultimately providing a window into the ancient marine biosphere (Konhauser et al., 2017). For example, a decline in nickel content in BIFs was used to hypothesize a “nickel famine” at ∼2.7 Ga in which biogenic methane production waned, allowing for the subsequent rise in atmospheric oxygenation at around 2.45 Ga, *i*.*e*., the Great Oxidation Event (GOE; Konhauser et al., 2009, 2015). In a similar manner, chromium enrichments in BIFs at 2.45 Ga were used to argue for enhanced oxidative weathering, most likely associated with the evolution of aerobic continental pyrite oxidation during the GOE (Konhauser et al., 2011). Increased oxidative weathering of land then not only supplied higher fluxes of critical nutrients (*e*.*g*., phosphorous; Bekker and Holland, 2012) but also toxic chemical species such as arsenate that would have added a new selective pressure on the survival of marine microbial communities (Chi Fru et al., 2019).

The utility of BIFs as a paleo-marine proxy for trace metals is predicated on the notion that the precipitation of Fe(III) oxyhydroxides, such as ferrihydrite (a likely precursor mineral to BIFs), captured trace metals in a predictable manner based on their partitioning coefficients in seawater. The ferrihydrite itself was precipitated in the photic zone overlying the continental shelf through the activity of marine phytoplankton, either directly by anoxygenic photosynthetic bacteria (the so-called photoferrotrophs) that use Fe(II) as their electron donor (Kappler and Pasquero, 2005; Posth et al., 2013) or indirectly via the abiogenic reaction of dissolved Fe^2+^ with O_2_ generated by primitive cyanobacteria (Cloud, 1965; Li et al., 2021). Upon phytoplankton death, some fraction of the cellular remains would have settled through the water column to the seafloor along with the biogenic ferrihydrite particles. In the sediment pile, the biomass would have undergone oxidation either during diagenesis via dissimilatory Fe(III)-reduction (DIR; Konhauser et al., 2005; Johnson et al., 2008) or during later metamorphism (Halama et al., 2016). Simultaneously, the ferrihydrite would have been reduced to dissolved Fe^2+^ that would have accumulated in the sediment porewaters and ultimately re-precipitated as an authigenic Fe phase, such as magnetite (Fe_3_O_4_), siderite (FeCO_3_) or some form of Fe-silicate phase (Li et al., 2013; Schad et al., 2021). The trace metals previously sorbed to the ferrihydrite would also have been solubilized and reincorporated into these authigenic mineral phases (Robbins et al., 2015).

This rather straightforward model, however, has generally glossed over the fact that the same phytoplankton that oxidized Fe(II) also contained a suite of trace metals in their biomass through assimilation for growth. Indeed, a recent modelling study suggested that photoferrotrophic biomass was a major contributor to trace elements (P, Mn, Co, Ni, Cu, Zn, Mo, Cd) incorporated into BIFs (Konhauser et al., 2018). Similar to the ferrihydrite, the decomposition of this microbial biomass in sediments by DIR bacteria would have resulted in the release of the trace elements originally associated with biomass back into pore waters where they were captured by Fe minerals. In summary, there likely were two major vectors for transferring metals from seawater into the sediment pile—sinking particles of ferrihydrite and dead plankton biomass—where the influence of DIR on the latter has not yet been experimentally confirmed.

In this work, we tested whether the degradation of biomass by a DIR bacterium liberated trace metals associated with the phytoplankton biomass. We incubated a model DIR bacterium (*Shewanella oneidensis* MR-1) under conditions that simulate Archean oceans (*i*.*e*., anoxic and silica-rich) with lysed phytoplankton biomass and ferrihydrite as a proxy for biogenic Fe(III) oxides. We collected subsamples for HCl-extractable Fe(II) and trace metal analyses to measure the reduction of Fe(III) and quantified changes in dissolved trace metal concentrations, respectively, as *S. oneidensis* MR-1 degrades the biomass and reduces the ferrihydrite. Although photoferrotrophs are presumed to be the major contributor of biomass and thus trace metals to BIFs prior to the GOE, cyanobacteria appeared as early as in the Paleoarchean (3.6–3.2 Ga; Sánchez-Baracaldo et al., 2021), and were another potential source of biomass to BIFs (Konhauser et al., 2018). Thus, we carried out parallel incubations using both photoferrotrophs and cyanobacteria as a source of phytoplankton biomass. Our results show that *S. oneidensis* MR-1 reduced Fe(II) when incubated with both photoferrotroph and cyanobacteria biomass, implying that this DIR bacteria is capable of consuming a small fraction of the carbon substrates from these biomass sources. However, trace metals associated with the biomass were not liberated into solution during this process. Thus, our broad interpretation is that the degradation of phytoplankton biomass by DIR bacteria would not have led to the release of trace metals into sediment porewaters where they could have potentially been captured by Fe minerals in the sediment and ultimately preserved in BIFs. Instead, other heterotrophic microbes would have been needed to degrade the metal-bearing compounds in phytoplankton biomass prior to the capture of these metals in precursor BIF sediments.

## 2. Methods

### 2.1 Dissimilatory Fe(III)-Reducing (DIR) Bacterium and Culture Conditions

*Shewanella oneidensis* MR-1 was used as a model DIR bacterial strain for this study. *S. oneidensis* MR-1 is facultative anaerobe with the capacity to reduce Fe(III) (oxyhydr)oxides (Lovley et al, 1989). *S. oneidensis* MR-1 was inoculated from a frozen glycerol stock into 15 mL of liquid Luria Bertani (LB) medium (Difco #244620) and incubated aerobically at 30°C while shaking at 140 rpm. When cultures reached an optical density at 600 nm (OD_600_) of 0.6 to 0.8 (mid-log phase; 6 h), 500 μL of culture was transferred to 50 mL of minimal (M1) medium (Kostka and Nealson, 1998) with 20 mM lactate as the electron donor. Cultures were grown aerobically at 30°C for 18 hours while shaking at 140 rpm to an OD_600_ of 0.8 to 0.9 (late-log phase) before incubation with phytoplankton biomass.

### 2.2 Growth medium and substrate synthesis

Experiments were conducted using M1 basal medium with the following modifications: amorphous silica (14.21 g L^-1^ Na_2_SiO_3_×9H_2_O) was added to simulate Archean seawater silica concentrations (final concentration: 2 mM; Maliva et al., 2005), and disodium anthraquinone-2,6 disulfonate (AQDS) was added to mediate the electron transfer between *S. oneidensis* MR-1 and ferrihydrite (final concentration 0.1 mM). The final modified M1 medium contained 3.3 mM KH_2_PO_4_, 5.7 mM K_2_HPO_4_, 9.9 μM NaCl, 45 μM H_3_BO_3_, 1.0 mM MgSO_4_×7H_2_O, 0.49 mM CaCl_2_×2H_2_O, 5.4 μM FeSO_4_×7H_2_O, 0.11 mM L-arginine, 0.19 mM L-serine, 0.14 mM L-glutamic acid, 11.5 μM Na_2_SeO_4_, 2.0 mM NaHCO_3_, 2 mM Na_2_SiO_3_×9H_2_O, and 0.1 mM AQDS.

Ferrihydrite was synthesized according to Schwertmann and Cornell (2000) using a 200 mM solution of Fe(NO_3_)_3_·9H_2_O neutralized with 1 M KOH to a final pH of 7.5. After centrifugation of the suspension for 10 min at 5000 g (Beckman Coulter Allegra X30-R Centrifuge), the supernatant was decanted and the wet solid was washed four times (resuspended in autoclaved MilliQ water, centrifuged, and decanted) to remove excess salts. Lastly, the wet solid was resuspended in 40 mL of autoclaved MilliQ water to a final concentration of 0.5 M. The formation of poorly crystalline 2-line ferrihydrite was confirmed by X-ray diffraction (XRD) using a Proto AXRD benchtop powder X-ray diffractometer at the University of Nevada, Las Vegas.

### 2.3 Preparation of Phytoplankton Biomass

Photoferrotroph (*Chlorobium ferroxidans* strain KoFox) and cyanobacteria (*Anabaena flos-aquae* and *Synechocystis sp*. PCC 6803) biomass served as carbon substrates during incubation of *S. oneidensis* MR-1. In the ocean, phytoplankton death leads to cell lysis and release of cellular contents, such that DIR in sediments would be primarily exposed to lysed biomass (*e.g*., Kirchman, 1999). Sonication is a commonly used laboratory method that uses sound waves to physically disrupt cell membranes and release cellular contents (*e.g*., proteins, nucleic acids, carbohydrates, etc.) for downstream analyses. Thus, photoferrotroph and cyanobacteria biomass were placed in an ultrasonic bath for 10 minutes to lyse cells (as confirmed via microscopic analysis) prior to incubation with ferrihydrite and *S. oneidensis* MR-1.

*Chlorobium ferrooxidans* strain KoFox was cultivated on modified freshwater medium containing 0.6 g L^-1^ KH_2_PO_4_, 0.3 g L^-1^ NH_4_Cl, 0.5 g L^-1^ MgSO_4_·7H_2_O and 0.1 g L^-1^ CaCl_2_·2H_2_O with a 22 mM bicarbonate buffer at pH 6.8-6.9 under an initial N_2_/CO_2_ headspace (90/10, *v/v*) as described in Hegler et al. (2008). The cultures were inoculated with 2% inoculum (*v/v*), the headspace exchanged for H_2_/CO_2_ (80/20, *v/v*) and the cultures incubated at 20°C in light (40 W incandescent light bulb) under static conditions. The headspace was exchanged every other day to consistently provide growth substrate (H_2_). Once the cultures reached stationary phase (determined by OD_600_) they were harvested by centrifugation (7000 g), washed three times with trace metal-free M1 medium to remove any trace metals adsorbed on cell walls, and freeze-dried.

The cyanobacteria biomass used in this experiment is comprised of 63% *Anabaena flos-aquae* and 27% *Synechocystis* sp. PCC 6803. Both cyanobacterial strains were grown in batch culture in liquid BG-11 medium (Rippka et al., 1979). Cultures were grown under cool white fluorescent lights, at 30°C, with gentle shaking to facilitate gas exchange and prevent cells from settling to the bottom of flasks. Cultures were grown under a 16:8 hour light:dark cycle, and culture density was monitored using optical density measured at 730 nm (OD_730_). When cultures reached late log phase, cells were harvested by centrifugation at 3000 g for 5 min. Cell pellets were washed three times by resuspending in growth media and centrifuging in between washes. Cell pellets were then stored at -80°, until freeze-drying prior to incorporation in experiments.

### 2.4 Anoxic Incubations with Phytoplankton Biomass

Modified M1 medium (12 mL) was aliquoted to sterile, acid-washed 25 mL glass serum bottles along with ferrihydrite (20 mM) and 12 mg of photoferrotroph or cyanobacteria biomass (42 mM organic carbon). Bottles were sealed with thick butyl rubber stoppers (Chemglass Life Sciences LLC; Vineland, NJ, USA) and crimped with aluminum seals. Subsequently, each bottle was purged with >99.998% N_2_ for 15 minutes. The serum bottles were then injected with 240 μL of *S. oneidensis* MR-1 log phase culture (2% inoculation, *v/v*) using a 22-gauge needle (BD PrecisionGlide™) and 1 mL syringe. All bottles were incubated in the dark at room temperature (22°C).

Several controls were included to track changes in Fe(III) reduction or trace metal concentrations in the absence of the *S. oneidensis* MR-1: (1) modified M1 medium, ferrihydrite, and photoferrotroph biomass; (2) modified M1 medium, ferrihydrite, cyanobacteria biomass; (3) modified M1 medium, ferrihydrite. In addition, controls with live *S. oneidensis* MR-1 in the absence of phytoplankton biomass were included to attribute any trends in Fe(III) reduction or trace metal concentrations to degradation of biomass: (1) modified M1 medium, ferrihydrite, 2% inoculation of *S. oneidensis* MR-1, and 10 mM lactate; (2) modified M1 medium, ferrihydrite, and a 2% inoculation of *S. oneidensis* MR-1 with no added carbon source. Experiments included two bottles of incubations of *S. oneidensis* MR-1 with ferrihydrite and photoferrotroph biomass, two bottles of incubations of *S. oneidensis* MR-1 with ferrihydrite and cyanobacteria biomass, and one bottle of each experimental control.

### 2.5 HCl-Extractable Fe(II)

In the presence of ferrihydrite, *S. oneidensis* MR-1 reduces the Fe(III) oxyhydroxide and produces Fe(II), making HCl-extractable Fe(II) a reliable proxy for the activity of this isolate (Lovley et al., 1989). Daily samples were taken for HCl-extractable Fe(II) from each bottle in replicate in an anoxic chamber (95% N_2_ and 5% H_2_; Coy Laboratory Products, Grass Lake, MI, USA) using a 23-gauge needle (BD PrecisionGlide™) and 1 mL plastic syringe. For HCl-extractable Fe(II) analyses (measured as dissolved Fe^2+^), 100 μL of shaken culture was extracted in replicate and placing it in 900 μL of 1 N HCl in the dark for 1 hr. Subsequently, the samples were centrifuged at 10000 g for 1 min, and 25 μL of supernatant was transferred into 975 μL of FerroZine reagent (buffered to pH 7.0 with 50 mM HEPES). Absorbance was measured within 5 minutes at 562 nm by UV-visible spectroscopy (Stookey, 1970). Iron concentrations were determined via comparison to a standard curve generated with ferrous ammonium sulfate. Reported values are an average of the two replicate measurements per bottle.

### 2.6 Inductively Coupled Plasma Mass Spectrometry (ICP-MS)

Changes in trace metal composition over the course of the experiment were evaluated via total dissolved trace metal concentrations. Bottles were sampled for trace metal analyses at the start (day 0), middle (day 5 for photoferrotroph biomass and day 6 for cyanobacteria biomass), and end (day 12) of each incubation. Samples were taken from each bottle in an anoxic chamber (95% N_2_ and 5% H_2_; Grass Lake, MI, USA) using a 23-gauge needle (BD PrecisionGlide™) and acid-washed 1 mL plastic syringe. To measure dissolved trace metal contents, sub-samples were filtered with 0.2 μm nylon filters (Basix™). All samples were placed into 4% v/v high purity HNO_3_ (Optima; Fisher Scientific Company; Nepean, ON, CA) in acid-washed 15-mL polypropylene tubes. Trace metal contents were measured on a Thermo Scientific iCAP Q ICP-MS at McGill University using yttrium as an internal standard. The instrument was calibrated with multi-element standards, and results were verified against AQUA-1 standard (National Research Council). Precision (calculated as relative standard deviation) based on the repeated analysis of standards was 1–9%, and based on technical replicates of samples was 1–5%. Reported values for incubations of *S. oneidensis* MR-1 with ferrihydrite and either photoferrotroph or cyanobacteria biomass are an average of two replicate bottles.

Total trace metal concentrations of phytoplankton biomass were measured to constrain the starting pool of metals available to *S. oneidensis* MR-1. Approximately 12 mg of freeze-dried phytoplankton and cyanobacteria biomass were each digested in 69% v/v high purity HNO_3_ (Optima; Fisher Scientific Company; Nepean, ON, CA) in capped 10 mL Teflon vessels (Savillex Purillex®; Eden Prairie, MN, USA) overnight (>12 hours) at 120°C. The digestant was diluted into 2% v/v high purity HNO_3_ (Optima; Fisher Scientific Company; Nepean, ON, CA) in acid-washed 15-mL polypropylene tubes. For the photoferrotroph biomass, Mn, Co, Ni, Zn, and Mo were found to be above detection limit while all other metals were below the detection limit (Supplemental Table 1). For the cyanobacteria biomass incubations, Mn, Cu, Zn, and Mo were above detection limit (Supplemental Table 1).

While we took precautions to reduce the contribution of trace metals to our incubations (*i.e*., acid-washing glassware, removing trace metal solutions from the M1 medium), some of the materials (*e.g*., chemicals used to make M1 media) contained trace metals. To further constrain the background inputs of trace metals in our incubations, we measured trace metal concentrations in the modified M1 media and ferrihydrite. Modified M1 medium and 20 mM ferrihydrite were each diluted in 4% v/v high purity HNO_3_ (Optima; Fisher Scientific Company; Nepean, ON, CA) in acid-washed 15-mL polypropylene tubes. The modified M1 medium and ferrihydrite had measurable concentrations of Mn and Mo, with all other metals below detection limit (Supplemental Table 1); however, these concentrations were several orders of magnitude lower than the photoferrotroph biomass (Supplemental Table 1). In contrast, the ferrihydrite had similar Mo concentrations to the cyanobacteria biomass (141 ± 0.94 nmol L^-1^ and 203 ± 90 nmol L^-1^, respectively; Supplemental Table 1); therefore, we do not consider changes in Mo contents when evaluating incubations with cyanobacteria biomass.

## 3. Results

### 3.1 Dissimilatory Fe(III) Reduction

During the 15 days of incubation of ferrihydrite (20 mM) with *S. oneidensis* MR-1 and the photoferrotroph and cyanobacterial biomass, dissolved Fe^2+^ concentrations increased to an average of 1.1 ± 0.3 mM and 1.8 ± 0.6 mM, respectively (Figure 1). Importantly, DIR with phytoplankton biomass is significantly less than in incubations of *S. oneidensis* MR-1 and lactate (6.6 ± 0.2 mM maximum; Supplemental Figure 1), implying that Fe(III) reduction is limited by the type of carbon source. Minimal increases in Fe^2+^ (< 0.3 mM) were measured for the abiotic controls containing modified M1 medium, ferrihydrite, and either photoferrotroph or cyanobacteria biomass. Evidence of abiotic Fe(III) reduction in our experiments is not surprising given that previous work has demonstrated that organic ligands—especially humic substances— can serve as electron donors in the reduction of Fe(III) (Kappler et al., 2021). While it remains unclear which organic ligands in the phytoplankton biomass were capable of reducing ferrihydrite, several organic molecules found in biomass, such as amino acids, have been shown to reduce Fe(III) (Bhattacharyya et al., 2019). Regardless, our data imply that the increased Fe^2+^ observed during incubation of *S. oneidensis* MR-1 and biomass is primarily attributable to dissimilatory Fe(III) reduction, and hence the metabolic activity of *S. oneidensis* MR-1.

**Figure 1.**
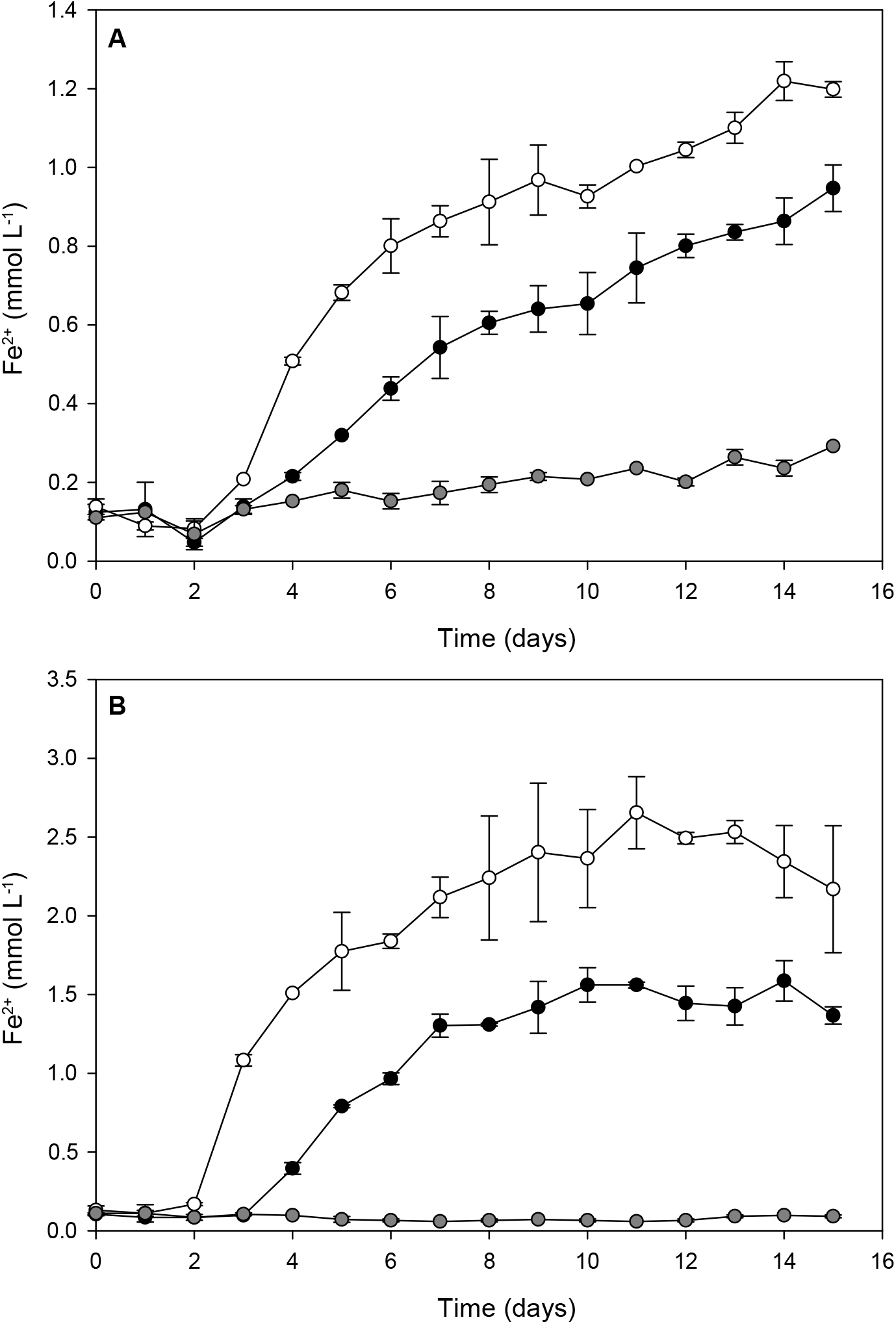
Reduction of ferrihydrite (20 mM) to Fe^2+^ by *S. oneidensis* MR-1 during incubation with A) photoferrotroph biomass (ca. 42 mM of organic carbon) and B) cyanobacteria biomass (ca. 42 mM of organic carbon) as a carbon source. Black and white symbols represent biotic replicates. Gray symbols represent abiotic controls with modified M1 medium, ferrihydrite and biomass. Data shown are the mean from duplicate measurements ± 1 standard deviation; bars not visible are smaller than the symbols.

### 3.2 Trace Metal Composition

Although we were successful in eliminating most trace metals from background sources (*e.g*., M1 media, glassware, etc.), we anticipated that the *S. oneidensis* MR-1 cells themselves would introduce trace metals to the background pool. Thus, we measured trace metal concentrations in live controls with modified M1 medium, ferrihydrite, and a 2% inoculation of *S. oneidensis* MR-1 with no added carbon source (Figures 2 and 3). By comparing these controls to our established background of trace metals in the modified M1 medium and ferrihydrite (Supplemental Table 1), we were able to confirm that some dissolved metals (Co, Ni, Zn, and Cu) were introduced by the inoculation of *S. oneidensis* MR-1. Therefore, trends in trace metal concentrations during incubation of *S. oneidensis* MR-1 with phytoplankton biomass were carefully considered in the context of the live control containing a 2% inoculation of *S. oneidensis* MR-1 (Figures 2 and 3; patterned bars) and abiotic controls containing modified M1 medium, ferrihydrite, and biomass (Figures 2 and 3; gray bars).

**Figure 2.**
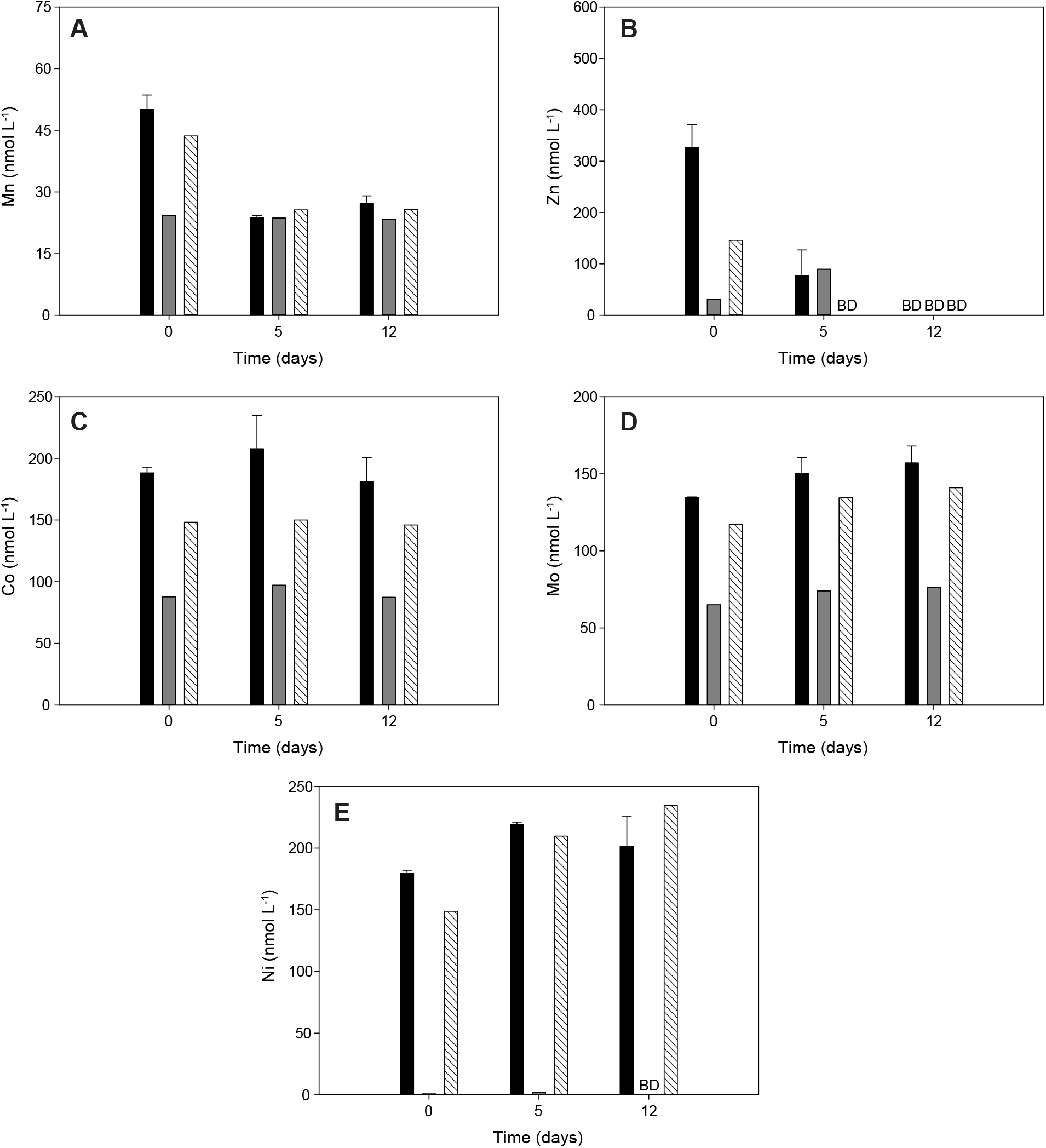
Dissolved concentrations of (A) manganese, (B) zinc, (C) cobalt, (D) molybdenum, and (E) nickel for days 0, 5, and 12 during incubation of *S. oneidensis* MR-1 with photoferrotroph biomass, modified M1 medium and ferrihydrite (black bars). Data shown include an abiotic control containing modified M1 medium, ferrihydrite and photoferrotroph biomass (gray bars), and a live control containing modified M1 medium, ferrihydrite and a 2% inoculation of *S. oneidensis* MR-1 (patterned bars). For the incubations with photoferrotroph biomass and *S. oneidensis* MR-1, data are the mean from duplicate measurements ± 1 standard deviation. Data marked “BD” are below the limit of detection.

**Figure 3.**
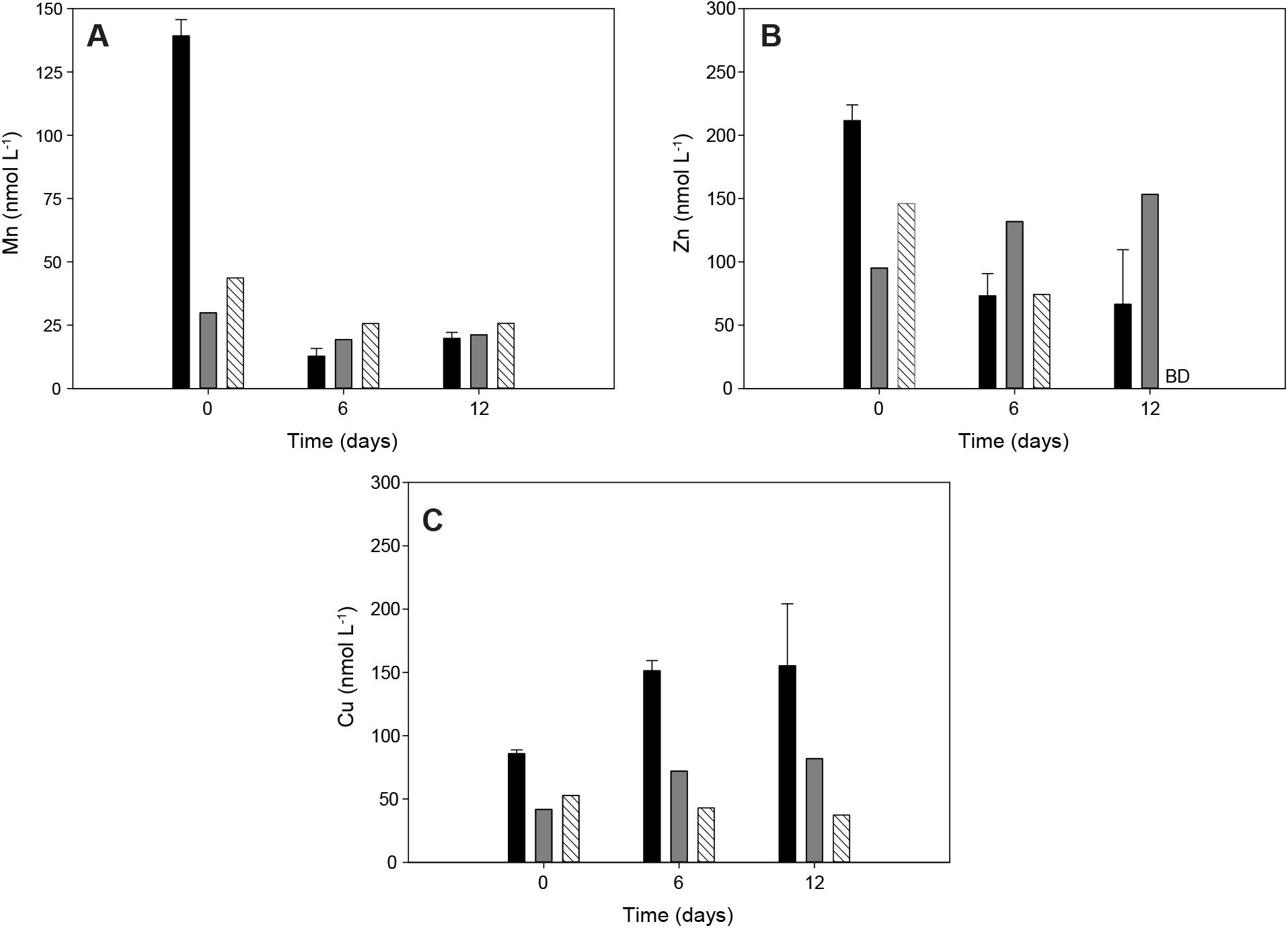
Dissolved concentrations of (A) manganese, (B) zinc, and (C) copper for days 0, 6, and 12 during incubation of *S. oneidensis* MR-1 with cyanobacteria biomass, modified M1 medium and ferrihydrite (black bars). Data shown include an abiotic control containing modified M1 medium, ferrihydrite and cyanobacteria biomass (gray bars), and a live control containing modified M1 medium, ferrihydrite and a 2% inoculation of *S. oneidensis* MR-1 (patterned bars). For the incubations with cyanobacteria biomass and *S. oneidensis* MR-1, data are the mean from duplicate measurements ± 1 standard deviation. Data marked “BD” are below the limit of detection. Note: the midpoint trace metal measurement for the live control (day 5) differed from the incubations with cyanobacteria biomass (day 6).

Characterization of the trace metals associated with the photoferrotroph biomass revealed that Mn, Co, Ni, Zn, and Mo were above detection limit (Supplemental Table 1), indicating that these metals were enriched in the biomass and could be potentially released during degradation of the biomass. For Mn and Zn, dissolved trace metal concentrations decreased substantially during incubation of *S. oneidensis* MR-1 with photoferrotroph biomass, suggesting no release of these metals over time. Specifically, dissolved Mn decreased by 23 ± 4 nmol L^-1^ (46% decrease; Figure 2A), and dissolved Zn decreased by 340 ± 65 nmol L^-1^ from days 0 to 12 (100% decrease; Figure 2B). For the other metals, such as dissolved Co (Figure 2C) and Mo (Figure 2D), their concentrations did not change over time and were consistent during incubation of *S. oneidensis* MR-1 with photoferrotroph biomass, as well as the both the abiotic and live controls. Lastly, although a slight increase (by 39.5 ± 0.5 nmol L^-1^; 22% increase) in dissolved Ni concentrations was observed from day 0 to 5 in bottles with *S. oneidensis* MR-1 and photoferrotroph biomass, increases were also observed in the live control (61 nmol L^-1^; 41% increase; Figure 2E), indicating that this increase is likely due to release of this metal from the *S. oneidensis* MR-1 cells as opposed to the degradation of biomass. Although there are no papers that have investigated the intracellular Ni concentrations of *S. oneidensis* MR-1, it is widely known that Ni-containing metalloproteins are often required for anaerobic microbial metabolisms (*e.g*., Alfano and Cavazza, 2020). Therefore, a pool of intracellular Ni associated with *S. oneidensis* MR-1 cells is a plausible source of the observed increase in dissolved Ni concentrations.

For the cyanobacteria biomass, trace metal analyses showed that Mn, Cu, Zn, and Mo were above detection limit and could be released from the biomass. However, Mo concentrations were low in the cyanobacterial biomass and comparable to background concentrations of Mo (Supplemental Table 1), thus changes in Mo were not considered during incubations with cyanobacteria biomass. Similar to incubations with photoferrotroph biomass, decreases in dissolved Mn and Zn concentrations were observed during incubation of *S. oneidensis* MR-1 with cyanobacteria biomass. Specifically, dissolved Mn and Zn concentrations decreased in bottles with *S. oneidensis* MR-1 and cyanobacteria biomass by 120 ± 16 nmol L^-^ (86% decrease; Figure 3A) and 145 ± 26 nmol L^-1^ (69% decrease; Figure 3B), respectively, over the course of the incubation. By contrast, an increase in dissolved Cu (by 70 ± 26 nmol L^-1^, 82% increase) was observed from days 0 to 12 during incubation of *S. oneidensis* MR-1 and cyanobacteria biomass (Figure 3C; black bars). However, this trend mirrors that of the abiotic control (modified M1 media, ferrihydrite and cyanobacteria biomass), in which dissolved Cu concentrations also increased by 96% (by 40 nmol L^-1^; Figure 3C; gray bars). Altogether, trends in trace metal concentrations between incubations of *S. oneidensis* MR-1 with phytoplankton biomass and experimental controls suggest no discernible trend in trace metal concentration that can be attributed to the degradation of biomass by *S. oneidensis* MR-1.

## 4. Discussion

In Archean oceans, an estimated 30% of phytoplankton biomass from the photic zone was buried in BIF precursor sediments (Konhauser et al., 2005). This organic material would have been a source of energy and carbon for microbes in ancient seafloor sediments and would have driven the transformation of ferrihydrite into minerals such as magnetite and siderite (*e.g*., Johnson et al. 2008; Konhauser et al., 2017), both major minerals in BIFs. In addition, if we assume that phytoplankton biomass served as a trace metal shuttle to sediments, microbial remineralization of biomass with ferrihydrite as the electron acceptor would have released significant quantities of trace metals back into the porewaters, eventually leading to the sequestration of these metals in precursor BIF sediments (Konhauser et al., 2018).

The Fe(III) reduction observed during our incubation experiments with phytoplankton biomass indicates that *S. oneidensis* MR-1 was able to access and consume carbon substrates needed for cellular growth and activity from both photoferrotroph and cyanobacteria biomass. However, Fe(III) reduction observed in incubations of *S. oneidensis* MR-1 with phytoplankton biomass was limited in comparison to controls with lactate. This limitation cannot be attributable to the amount of ferrihydrite present in all incubations (20 mM), nor the amount of organic carbon present (30 mM in the lactate experiments, 42 mM in the incubations with biomass).

Instead, the Fe(III) reduction in our experiments must have been limited by the bioavailability of organic macromolecules present in the phytoplankton biomass. A recent estimate compiled from 222 marine and freshwater species found that the median macromolecular composition of phytoplankton (based on dry weight) is 32% protein, 17% lipids, 15% carbohydrates, 7% nucleic acids (Finkel et al., 2016). In our incubation experiments, phytoplankton biomass was sonicated which would have lysed the cells and released these biomolecules into the surrounding medium.

Microbes generally cannot take up these macromolecules directly, so they rely on extracellular degradative enzymes to break down these molecules to less than ∼600 Da prior to uptake into the cytoplasm (Arnosti, 2011). The genome for *S. oneidensis* MR-1 has been well studied and does not encode for any known extracellular peptidases that would be required to initiate the degradation of metalloproteins (Heidelberg et al., 2002). Likewise, *S. oneidensis* MR-1 does not appear to have the metabolic capability to secrete glucosidases and lipases which would be needed to cleave complex carbohydrates and lipids, respectively, into oligomers and monomers. Interestingly, *S. oneidensis* MR-1 encodes for three different extracellular endonucleases that enable it to access nucleic acids (Gödeke et al., 2011; Heun et al., 2012). This is consistent with laboratory experiments which have demonstrated that *S oneidensis* MR-1 can consume extracellular DNA as a sole carbon and energy source under anoxic conditions (Pinchuk et al., 2008). Additional laboratory studies have shown that *S. oneidensis* MR-1 consumes simple substrates such as pyruvate (Lovley et al., 1989) and monosaccharides (Hunt et al., 2010) — both of which are found in phytoplankton cells. Although we cannot identify the specific biomolecules, it is likely that *S. oneidensis* MR-1 consumed a combination of some biomolecules as a source of carbon during our incubations. Taken together, these observations are consistent with other well-known DIR bacteria found in marine sediments (*e.g*., *Geobacter metallireducens*) that do not appear to secrete degradative enzymes and instead rely on small carbon substrates (Lovley et al., 1997, and references therein).

We did not observe any accumulation of dissolved trace metal concentrations in the liquid phase as a function of time, which would provide evidence that *S. oneidensis* MR-1 was degrading the metalloproteins associated with phytoplankton biomass, and subsequently liberating these trace metals. Additionally, there does not appear to be evidence of the incorporation of trace metals into *S. oneidensis* MR-1 cells during growth (*i.e*., a decrease in dissolved trace metal content that differs from the abiotic and live controls), implying that other mechanisms caused the trends in trace metal composition in our experiments. Interestingly, trace metal concentrations in our incubations with abiotic controls did not remain constant over time (*e.g*., Zn and Cu; incubations with cyanobacteria biomass), indicating that other processes or interactions can influence the partitioning of trace metals between dissolved and particulate (*e.g*., ferrihydrite) fractions and would ultimately impact how trace metals are incorporated into precursor BIF sediments. Previous work examining the release of trace metals during phytoplankton decay has demonstrated that metals have different affinities for retention in biomass (*e.g*., Co is more conservative than Zn; Hollister et al., 2020). The presence of ferrihydrite and silica in our incubations complicates our interpretations of trace metal trends even further because Fe(III) oxyhydroxides like ferrihydrite can strongly bind trace metals, albeit with different binding affinities for different metals (Kappler et al., 2021). Adsorption of trace metals to ferrihydrite may be stunted somewhat by amorphous silica, which binds to adsorption sites on the ferrihydrite and can outcompete other ions like phosphate, and, potentially, trace metals (Konhauser et al., 2009). Organic matter (*e.g*., phytoplankton biomass) would also affect the extent to which metals are adsorbed to ferrihydrite, because the organic matter simultaneously competes for metal adsorption sites on the ferrihydrite while also providing additional binding sites itself (Engel et al., 2021). Thus, trace metal adsorption to ferrihydrite and dissolved phases differs when comparing metal sorption to ferrihydrite alone and metal sorption to ferrihydrite in the presence of biomass, with some metals preferentially associated with ferrihydrite (*e.g*., Ni; Moon and Peacock, 2013), and other metals having a strong affinity for biomass (*e.g*., Cu; Eickhoff et al., 2014). Lastly, the reduction of ferrihydrite by DIR bacteria could lead to secondary mineral formation (to magnetite, goethite, and hematite) even over the course of a few days (Xiao et al., 2018). This, in turn, would subsequently impact the adsorption and release of trace metals from biomass and Fe minerals. Although the trace metal concentrations evaluated in this work do not provide enough resolution to identify the extent of trace metal release and adsorption to ferrihydrite, these adsorption pathways must be considered when evaluating how biogenic trace metals are incorporated into BIFs.

Our study provides a framework from which to evaluate the role of DIR in the recycling of phytoplankton biomass in precursor BIF sediments. The results strongly point to additional heterotrophic microorganisms as playing a key role in the release of trace metals associated with phytoplankton biomass back into sediment porewaters for their eventual preservation in BIFs. In modern anoxic marine sediments, organic matter is degraded through a sequence of steps with different microorganisms involved (Arndt et al., 2013). The first step is the extracellular degradation of complex macromolecules by primary degraders followed by the fermentation of oligomers and monomers to alcohols, lactate and volatile fatty acids, which are then mineralized to CH_4_, CO_2_ and/or H_2_. It remains unclear as to whether primary degraders and fermenters are in fact two separate populations or if fermenters are both secreting degradative enzymes and fermenting the resulting degradation products. The metabolic capability to produce extracellular enzymes is widespread across taxa (Zimmerman et al., 2013; Arnosti, 2011) and genomic reconstructions have revealed the presence of these enzymes in non-fermenting microbes (Lloyd et al., 2013). More recently, a study that investigated the anerobic degradation of organic matter in marine sediments using ^13^C-labeled proteins and lipids found primary degraders that also encoded pathways to ferment the degradation products such as amino acids (Pelikan et al., 2020). Given that trace metals within cells are typically associated with complex proteins (*e.g*., metalloproteins; Jelen et al., 2016), the importance of primary degraders cannot be overstated as they would have initiated the remineralization of metal-bearing compounds in phytoplankton biomass which have led to the release of trace metals back into the water column or sediment porewaters. Aside from DIR bacteria, methanogens and fermenters are thought to have been present in ancient Archaean sediments (Konhauser et al., 2005; Posth et al., 2013). Regardless of whether fermenters were the primary group to initiate the degradation of macromolecules in Archaean sediments, DIR bacteria would likely rely on fermenters to produce small compounds that could be readily assimilated as a carbon and energy such that the activity DIR bacteria would be limited in the absence of fermenters. Future studies focused on resolving the role and function of other heterotrophic microbes in the remineralization of phytoplankton biomass are crucial to advancing our understanding of BIFs and their function as an ancient recorder of seawater chemistry.

## Acknowledgements

This work was supported financially by a Wares Postdoctoral Fellowship to KR and a Natural Sciences & Engineering Research Council of Canada (NSERC) grant to NM. AK acknowledges infrastructural support by the Deutsche Forschungsgemeinschaft (DFG, German Research Foundation) under Germany’s Excellence Strategy, cluster of Excellence EXC2124, project ID 390838134. We thank Ana Gonzalez-Nayeck (Harvard University) and Jenan Kharbush (University of Michigan) for providing cyanobacteria biomass, as well as Anna Jung (McGill) for ICP analysis and Thi Hao Bui (McGill) for technical support with anoxic incubations. We would also like to thank Jennifer Glass (Georgia Institute of Technology) and Elliott Mueller (California Institute of Technology) for thoughtful discussions that improved this study.

---

**Supplemental Figure 1.**
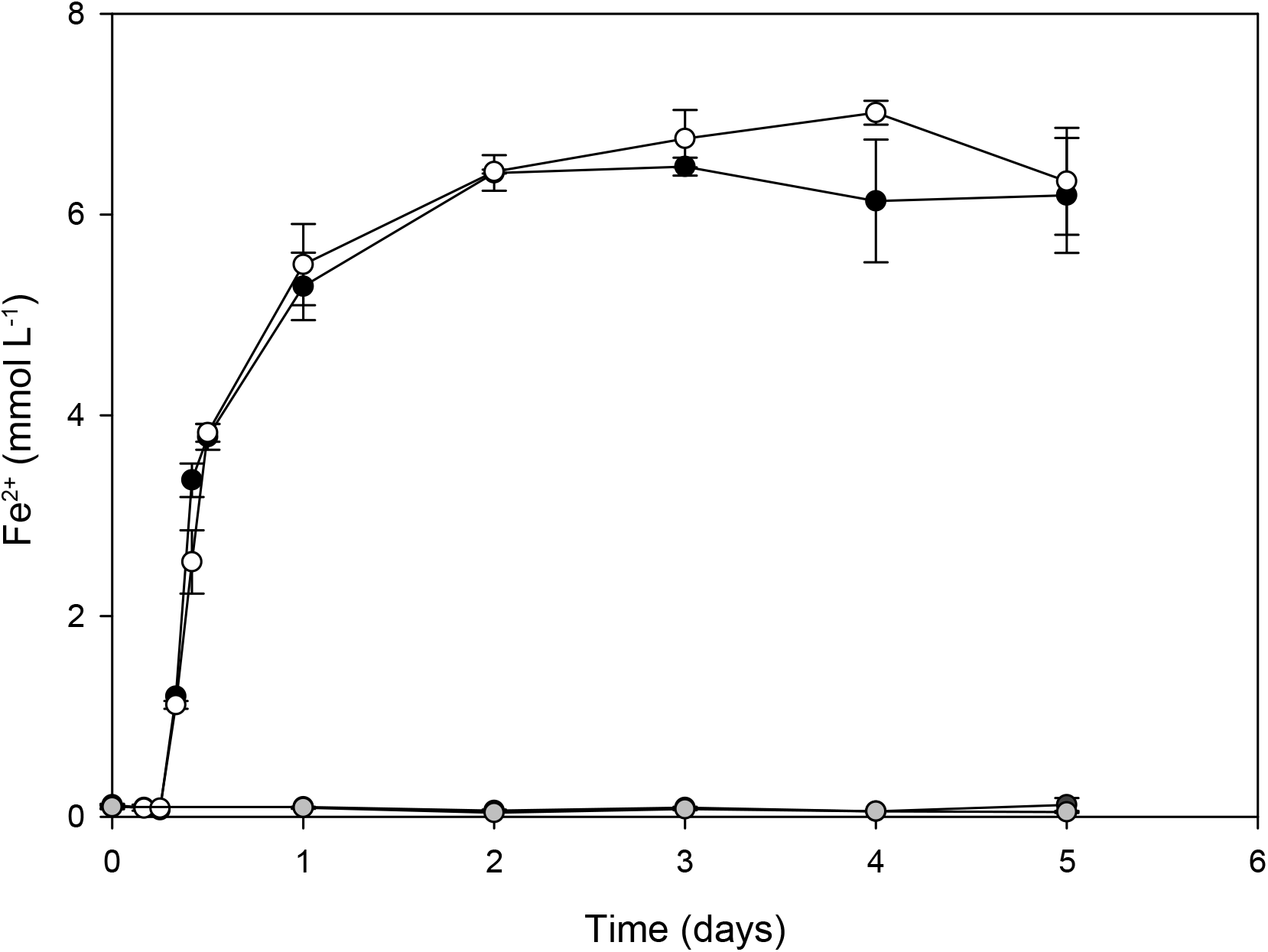
Reduction of ferrihydrite to Fe^2+^ by *S. oneidensis* MR-1 during incubation with 20 mM ferrihydrite and 10 mM lactate as the electron donor. Black and white symbols represent biotic replicates. Dark gray symbols represent abiotic controls containing modified M1 medium and ferrihydrite, while light gray symbols represent live controls containing a modified M1 medium, ferrihydrite and a 2% inoculation of *S. oneidensis* MR-1; data overlap and thus abiotic control data is not easily visible. Data shown are the mean from duplicate measurements ± 1 standard deviation; bars not visible are smaller than the symbols.

**Supplemental Table 1.** Trace metal composition of the modified M1 medium, ferrihydrite, photoferrotroph biomass, and cyanobacteria biomass. For all materials, Cd concentrations were below detection limit. Data marked “*B.D*.” are below the limit of detection.

|  | Modified M1<br>Medium | Ferrihydrite | Photoferrotroph<br>Biomass | Cyanobacteria<br>Biomass |
| --- | --- | --- | --- | --- |
| Mn (nmol L<sup>-1</sup>) | 40 ± 3 | 407 ± 5 | 28733 ± 360 | 5435137 ± 23153 |
| Co (nmol L<sup>-1</sup>) | *B.D.* | *B.D.* | 1285474 ± 10476 | *B.D.* |
| Ni (nmol L<sup>-1</sup>) | *B.D.* | *B.D.* | 24186 ± 1607 | *B.D.* |
| Cu (nmol L<sup>-1</sup>) | *B.D.* | *B.D.* | *B.D.* | 8984 ± 1136 |
| Zn (nmol L<sup>-1</sup>) | *B.D.* | *B.D.* | 1108645 ± 4251 | 824696 ± 2837 |
| Mo (nmol L<sup>-1</sup>) | 27 ± 7 | 141 ± 1 | 27769 ± 111 | 203 ± 90 |

## References

Alfano, M., Cavazza, C., 2020. Structure, function, and biosynthesis of nickel-dependent enzymes. Protein Sci. 29, 1071–1089. https://doi.org/10.1002/pro.3836

Arndt, S., Jørgensen, B.B., LaRowe, D.E., Middelburg, J.J., Pancost, R.D., Regnier, P., 2013. Quantifying the degradation of organic matter in marine sediments: A review and synthesis. Earth-Science Rev. 123, 53–86. https://doi.org/10.1016/j.earscirev.2013.02.008

Arnosti, C., 2011. Microbial extracellular enzymes and the marine carbon cycle. Ann. Rev. Mar. Sci. 3, 401–425. https://doi.org/10.1146/annurev-marine-120709-142731

Bekker, A., Holland, H.D., 2012. Oxygen overshoot and recovery during the early Paleoproterozoic. Earth Planet. Sci. Lett. 317–318, 295–304. https://doi.org/10.1016/j.epsl.2011.12.012

Bhattacharyya, A., Schmidt, M.P., Stavitski, E., Azimzadeh, B., Martínez, C.E., 2019. Ligands representing important functional groups of natural organic matter facilitate Fe redox transformations and resulting binding environments. Geochim. Cosmochim. Acta 251, 157– 175. https://doi.org/10.1016/j.gca.2019.02.027

Chi Fru, E., Somogyi, A., El Albani, A., Medjoubi, K., Aubineau, J., Robbins, L.J., Lalonde, S. V., Konhauser, K.O., 2019. The rise of oxygen-driven arsenic cycling at ca. 2.48 Ga. Geology 47, 243–246. https://doi.org/10.1130/G45676.1

Cloud, P.E., 1965. Significance of the Gunflint (Precambrian) Microflora: Photosynthetic oxygen may have had important local effects before becoming a major atmospheric gas. Science (80-.). 148, 27–35. https://doi.org/10.1126/science.148.3666.27

Eickhoff, M., Obst, M., Schröder, C., Hitchcock, A.P., Tyliszczak, T., Martinez, R.E., Robbins, L.J., Konhauser, K.O., Kappler, A., 2014. Nickel partitioning in biogenic and abiogenic ferrihydrite: The influence of silica and implications for ancient environments. Geochim. Cosmochim. Acta 140, 65–79. https://doi.org/10.1016/j.gca.2014.05.021

Engel, M., Lezama Pacheco, J.S., Noël, V., Boye, K., Fendorf, S., 2021. Organic compounds alter the preference and rates of heavy metal adsorption on ferrihydrite. Sci. Total Environ. 750. https://doi.org/10.1016/j.scitotenv.2020.141485

Finkel, Z. V., Follows, M.J., Liefer, J.D., Brown, C.M., Benner, I., Irwin, A.J., 2016. Phylogenetic diversity in the macromolecular composition of microalgae. PLoS One 11, 1– 16. https://doi.org/10.1371/journal.pone.0155977

Gödeke, J., Heun, M., Bubendorfer, S., Paul, K., Thormann, K.M., 2011. Roles of two Shewanella oneidensis MR-1 extracellular endonucleases. Appl. Environ. Microbiol. 77, 5342–5351. https://doi.org/10.1128/AEM.00643-11

Halama, M., Swanner, E.D., Konhauser, K.O., Kappler, A., 2016. Evaluation of siderite and magnetite formation in BIFs by pressure–temperature experiments of Fe(III) minerals and microbial biomass. Earth Planet. Sci. Lett. 450, 243–253. https://doi.org/10.1016/j.epsl.2016.06.032

Hegler, F., Posth, N.R., Jiang, J., Kappler, A., 2008. Physiology of phototrophic iron(II)-oxidizing bacteria: Implications for modern and ancient environments. FEMS Microbiol. Ecol. 66, 250–260. https://doi.org/10.1111/j.1574-6941.2008.00592.x

Heidelberg, J.F., Paulsen, I.T., Nelson, K.E., Gaidos, E.J., Nelson, W.C., Read, T.D., Eisen, J.A., Seshadri, R., Ward, N., Methe, B., Clayton, R.A., Meyer, T., Tsapin, A., Scott, J., Beanan, M., Brinkac, L., Daugherty, S., DeBoy, R.T., Dodson, R.J., Durkin, A.S., Haft, D.H., Kolonay, J.F., Madupu, R., Peterson, J.D., Umayam, L.A., White, O., Wolf, A.M., Vamathevan, J., Weidman, J., Impraim, M., Lee, K., Berry, K., Lee, C., Mueller, J., Khouri, H., Gill, J., Utterback, T.R., McDonald, L.A., Feldblyum, T. V., Smith, H.O., Venter, J.C., Nealson, K.H., Fraser, C.M., 2002. Genome sequence of the dissimilatory metal ion-reducing bacterium Shewanella oneidensis. Nat. Biotechnol. 20, 1118–1123. https://doi.org/10.1038/nbt749

Heun, M., Binnenkade, L., Kreienbaum, M., Thormann, K.M., 2012. Functional specificity of extracellular nucleases of Shewanella oneidensis MR-1. Appl. Environ. Microbiol. 78, 4400–4411. https://doi.org/10.1128/AEM.07895-11

Hollister, A.P., Kerr, M., Malki, K., Muhlbach, E., Robert, M., Tilney, C.L., Breitbart, M., Hubbard, K.A., Buck, K.N., 2020. Regeneration of macronutrients and trace metals during phytoplankton decay: An experimental study. Limnol. Oceanogr. 1936–1960. https://doi.org/10.1002/lno.11429

Hunt, K.A., Flynn, J.M., Naranjo, B., Shikhare, I.D., Gralnick, J.A., 2010. Substrate-level phosphorylation is the primary source of energy conservation during anaerobic respiration of Shewanella oneidensis strain MR-1. J. Bacteriol. 192, 3345–3351. https://doi.org/10.1128/JB.00090-10

Jelen, B.I., Giovannelli, D., Falkowski, P.G., 2016. The Role of Microbial Electron Transfer in the Coevolution of the Biosphere and Geosphere. Annu. Rev. Microbiol. 70, 45–62. https://doi.org/10.1146/annurev-micro-102215-095521

Johnson, C.M., Beard, B.L., Roden, E.E., 2008. The iron isotope fingerprints of redox and biogeochemical cycling in modern and ancient earth. Annu. Rev. Earth Planet. Sci. 36, 457– 493. https://doi.org/10.1146/annurev.earth.36.031207.124139

Kappler, A., Bryce, C., Mansor, M., Lueder, U., Byrne, J.M., Swanner, E.D., 2021. An evolving view on biogeochemical cycling of iron. Nat. Rev. Microbiol. https://doi.org/10.1038/s41579-020-00502-7

Kappler, A., Pasquero, C., 2005. Deposition of banded iron formations by anoxygenic phototrophic Fe (II) -oxidizing bacteria. Geology 865–868. https://doi.org/10.1130/G21658.1

Kirchman, D.L., 1999. Phytoplankton Death in the Sea. Nat. News views 398, 293–294.

Konhauser, K.O., Planavsky, N.J., Hardisty, D.S., Robbins, L.J., Warchola, T.J., Haugaard, R., Lalonde, S. V., Partin, C.A., Oonk, P.B.H., Tsikos, H., Lyons, T.W., Bekker, A., Johnson, C.M., 2017. Iron formations: A global record of Neoarchaean to Palaeoproterozoic environmental history. Earth-Science Rev. 172, 140–177. https://doi.org/10.1016/j.earscirev.2017.06.012

Konhauser, K.O., Newman, D.., Kappler, A., 2005. The potential significance of microbial Fe(III) reduction during deposition of Precambrian banded iron formations. Geobiology 3, 167–177.

Konhauser, K.O., Lalonde, S. V., Planavsky, N.J., Pecoits, E., Lyons, T.W., Mojzsis, S.J., Rouxel, O.J., Barley, M.E., Rosìere, C., Fralick, P.W., Kump, L.R., Bekker, A., 2011. Aerobic bacterial pyrite oxidation and acid rock drainage during the Great Oxidation Event. Nature 478, 369–373. https://doi.org/10.1038/nature10511

Konhauser, K.O., Pecoits, E., Lalonde, S. V., Papineau, D., Nisbet, E.G., Barley, M.E., Arndt, N.T., Zahnle, K., Kamber, B.S., 2009. Oceanic nickel depletion and a methanogen famine before the Great Oxidation Event. Nature 458, 750–753. https://doi.org/10.1038/nature07858

Konhauser, K.O., Robbins, L.J., Alessi, D.S., Flynn, S.L., Gingras, M.K., Martinez, R.E., Kappler, A., Swanner, E.D., Li, Y.L., Crowe, S.A., Planavsky, N.J., Reinhard, C.T., Lalonde, S. V., 2018. Phytoplankton contributions to the trace-element composition of Precambrian banded iron formations. Bull. Geol. Soc. Am. 130, 941–951. https://doi.org/10.1130/B31648.1

Konhauser, K.O., Robbins, L.J., Pecoits, E., Peacock, C., Kappler, A., Lalonde, S. V., 2015. The Archean Nickel Famine Revisited. Astrobiology 15, 804–815. https://doi.org/10.1089/ast.2015.1301

Kostka, J.E., Nealson, K.H., 1998. Isolation, cultivation, and characterization of iron- and manganese-reducing bacteria, in: Burlage, R.S. (Ed.), Techniques in Microbial Ecology. Oxford University Press, Oxford, UK, pp. 58–78.

Li, Y.L., Konhauser, K.O., Kappler, A., Hao, X.L., 2013. Experimental low-grade alteration of biogenic magnetite indicates microbial involvement in generation of banded iron formations. Earth Planet. Sci. Lett. 361, 229–237. https://doi.org/10.1016/j.epsl.2012.10.025

Li, Y., Sutherland, B.R., Gingras, M.K., Owttrim, G.W., Konhauser, K.O., 2021. A novel approach to investigate the deposition of (bio)chemical sediments: The sedimentation velocity of cyanobacteria-ferrihydrite aggregates. J. Sediment. Res. 91, 390–398. https://doi.org/10.2110/JSR.2020.114

Lloyd, K.G., Schreiber, L., Petersen, D.G., Kjeldsen, K.U., Lever, M.A., Steen, A.D., Stepanauskas, R., Richter, M., Kleindienst, S., Lenk, S., Schramm, A., Jorgensen, B.B., 2013. Predominant archaea in marine sediments degrade detrital proteins. Nature 496, 215– 218. https://doi.org/10.1038/nature12033

Lovley, D.R., Phillips, E.J.P., Lonergan, D.J., 1989. Hydrogen and formate oxidation coupled to dissimilatory reduction of iron or manganese by Alteromonas putrefaciens. Appl. Environ. Microbiol. 55, 700–706. https://doi.org/10.1128/aem.55.3.700-706.1989

Lovley, D.R., Coates, J.D., Saffarini, D.A., Lonergan, D.J., 1997. Dissimilatory Iron Reduction, in: Winkelmann, G., Carrano, C.J. (Eds.), Transition Metals in Microbial Metabolism. hardwood academic publishers, Amsterdam, pp. 187–215.

Lyons, T.W., Reinhard, C.T., Planavsky, N.J., 2014. The rise of oxygen in Earth’s early ocean and atmosphere. Nature 506, 307–15. https://doi.org/10.1038/nature13068

Maliva, R.G., Knoll, A.H., Simonson, B.M., 2005. Secular change in the Precambrian silica cycle: Insights from chert petrology. Bull. Geol. Soc. Am. 117, 835–845. https://doi.org/10.1130/B25555.1

Moon, E.M., Peacock, C.L., 2013. Modelling Cu(II) adsorption to ferrihydrite and ferrihydrite-bacteria composites: Deviation from additive adsorption in the composite sorption system. Geochim. Cosmochim. Acta 104, 148–164. https://doi.org/10.1016/j.gca.2012.11.030

Pelikan, C., Wasmund, K., Glombitza, C., Hausmann, B., Herbold, C.W., Flieder, M., Loy, A., 2021. Anaerobic bacterial degradation of protein and lipid macromolecules in subarctic marine sediment. ISME J. 15, 833–847. https://doi.org/10.1038/s41396-020-00817-6

Pinchuk, G.E., Ammons, C., Culley, D.E., Li, S.M.W., McLean, J.S., Romine, M.F., Nealson, K.H., Fredrickson, J.K., Beliaev, A.S., 2008. Utilization of DNA as a sole source of phosphorus, carbon, and energy by Shewanella spp.: Ecological and physiological implications for dissimilatory metal reduction. Appl. Environ. Microbiol. 74, 1198–1208. https://doi.org/10.1128/AEM.02026-07

Posth, N.R., Konhauser, K.O., Kappler, A., 2013. Microbiological processes in banded iron formation deposition. Sedimentology 60, 1733–1754. https://doi.org/10.1111/sed.12051

Rippka, R., Deruelles, J., Waterbury, J.B., 1979. Generic assignments, strain histories and properties of pure cultures of cyanobacteria. J. Gen. Microbiol. 111, 1–61. https://doi.org/10.1099/00221287-111-1-1

Robbins, L.J., Lalonde, S. V., Planavsky, N.J., Partin, C.A., Reinhard, C.T., Kendall, B., Scott, C., Hardisty, D.S., Gill, B.C., Alessi, D.S., Dupont, C.L., Saito, M.A., Crowe, S.A., Poulton, S.W., Bekker, A., Lyons, T.W., Konhauser, K.O., 2016. Trace elements at the intersection of marine biological and geochemical evolution. Earth-Science Rev. 163, 323– 348. https://doi.org/10.1016/j.earscirev.2016.10.013

Robbins, L.J., Swanner, E.D., Lalonde, S. V., Eickhoff, M., Paranich, M.L., Reinhard, C.T., Peacock, C.L., Kappler, A., Konhauser, K.O., 2015. Limited Zn and Ni mobility during simulated iron formation diagenesis. Chem. Geol. 402, 30–39. https://doi.org/10.1016/j.chemgeo.2015.02.037

Sanchez-Baracaldo, P Bianchini, G., Wilson, J., Knoll, A., 2021. Cyanobacteria and biogeochemical cycles through Earth history. Trends Microbiol. In press, 1–15. https://doi.org/10.1016/j.tim.2021.05.008

Schad, M., Halama, M., Jakus, N., Robbins, L.J., Warchola, T.J., Tejada, J., Kirchhof, R., Lalonde, S. V., Swanner, E.D., Planavsky, N.J., Thorwarth, H., Mansor, M., Konhauser, K.O., Kappler, A., 2021. Phosphate remobilization from banded iron formations during metamorphic mineral transformations. Chem. Geol. 584. https://doi.org/10.1016/j.chemgeo.2021.120489

Schwertmann, U., Cornell, R.M., 2000. Iron Oxides in the Laboratory: Preparation and Characterization. WILEY_VHC, Weinheim, Germany.

Stookey, L.L., 1970. Ferrozine-A New Spectrophotometric Reagent for Iron. Anal. Chem. 42, 779–781. https://doi.org/10.1021/ac60289a016

Swanner, E.D., Planavsky, N.J., Lalonde, S. V., Robbins, L.J., Bekker, A., Rouxel, O.J., Saito, M.A., Kappler, A., Mojzsis, S.J., Konhauser, K.O., 2014. Cobalt and marine redox evolution. Earth Planet. Sci. Lett. 390, 253–263. https://doi.org/10.1016/j.epsl.2014.01.001

Xiao, W., Jones, A.M., Li, X., Collins, R.N., Waite, T.D., 2018. Effect of Shewanella oneidensis on the Kinetics of Fe(II)-Catalyzed Transformation of Ferrihydrite to Crystalline Iron Oxides. Environ. Sci. Technol. 52, 114–123. https://doi.org/10.1021/acs.est.7b05098

Zimmerman, A.E., Martiny, A.C., Allison, S.D., 2013. Microdiversity of extracellular enzyme genes among sequenced prokaryotic genomes. ISME J. 7, 1187–1199. https://doi.org/10.1038/ismej.2012.176

